# Recurrent network model for learning goal-directed sequences through reverse replay

**DOI:** 10.1101/217422

**Authors:** Tatsuya Haga, Tomoki Fukai

## Abstract

Reverse replay of hippocampal place cells occurs frequently at rewarded locations, suggesting its contribution to goal-directed pathway learning. Symmetric spike-timing dependent plasticity (STDP) in CA3 likely potentiates recurrent synapses for both forward (start to goal) and reverse (goal to start) replays during sequential activation of place cells. However, how reverse replay selectively strengthens forward pathway is unclear. Here, we show computationally that firing sequences bias synaptic transmissions to the opposite direction of propagation under symmetric, but not asymmetric, STDP in the co-presence of short-term synaptic plasticity. We demonstrate that a significant bias can be created in biologically realistic simulation settings, and that this bias enables reverse replay to enhance goal-directed spatial memory on a T-maze. Our model for the first time provides the mechanistic account for the way reverse replay contributes to hippocampal sequence learning for reward-seeking spatial navigation.

## INTRODUCTION

In the rodent hippocampus, firing sequences of place cells are replayed during awake immobility and sleep (Carr et al., 2011). Replay can be either in the same firing order as experienced (forward replay) or in the reversed order (reverse replay). Forward replay is observed during sleep after exploration (Lee and Wilson, 2002) or in immobile states before rats start to travel towards reward (Diba and Buzsáki, 2007; Pfeiffer and Foster, 2013), hence forward replay is thought to engage in the consolidation and retrieval of spatial memory. In contrast, reverse replay presumably contributes to the optimization of goal-directed pathways because rewarded pathways are replayed around the timing of reward delivery (Diba and Buzsáki, 2007; Foster and Wilson, 2006), and the occurrence frequency is modulated by the presence and the amount of reward (Ambrose et al., 2016; Singer and Frank, 2009).

Because sequences are essentially time asymmetric, sequence learning often hypothesizes Hebbian spike-timing-dependent plasticity (STDP) with asymmetric time windows, which induce long-term potentiation (LTP) for pre-to-post firing order and long-term depression (LTD) for post-to-pre firing order. In the hippocampal area CA1 STDP is Hebbian (Bi and Poo, 2001) and is known to be enhanced by dopamine, which converts the LTD component into potentiation (Zhang et al., 2009). This modulation can occur even if dopamine is induced after the pairing of pre- and postsynaptic spikes (Brzosko et al., 2015), supporting the hypothesis that reward-induced signaling consolidates the synaptic eligibility trace established by preceding experience-dependent neuronal firing (Izhikevich, 2007). However, it was recently reported that the default form of STDP is time symmetric at recurrent synapses in the hippocampal area CA3 (Mishra et al., 2016). Because CA3 is the most likely source of replay sequences (Nakashiba et al., 2009), this finding raises the question whether and how STDP underlies sequence learning in CA3. A symmetric time window implies that reverse replay equally strengthens both forward synaptic pathways leading to the rewarded location and reverse pathways leaving away from the rewarded location in CA3 recurrent network. However, reward-based optimization requires selective reinforcement of forward pathways as it will strengthen prospective place-cell sequences in subsequent trials and forward replay events in the consolidation phase. How this directionality arises in replay events and how reverse replay enables the learning of goal-directed navigation remain unclear.

In this paper, we first show how goal-directed pathway learning is naturally realized through reverse replay in a one-dimensional chain model. To this end, we hypothesize that the contribution of presynaptic spiking for STDP is attenuated in CA3 by short-term depression, as was revealed in the rat visual cortex (Froemke et al., 2006). Under this condition, symmetric STDP and a rate-based Hebbian plasticity rule bias recurrent synaptic weights towards the opposite direction to the propagation of a firing sequence, implying that the combined rule virtually acts like anti-Hebbian STDP. By simulating the model with various input spike trains, we confirm this effect for a broad range of input spike trains and parameters of plasticity rules, including those observed in experiments. Finally, we demonstrate that our model works in a two-dimensional recurrent network of place cells to optimize forward pathways leading to reward by the combination of reverse replay, Hebbian plasticity with short-term plasticity, and dopaminergic enhancement of replay frequency (Ambrose et al., 2016; Singer and Frank, 2009). Unlike the previous models for hippocampal sequence learning (Blum and Abbott, 1996; Gerstner and Abbott, 1997; Jahnke et al., 2015; Jensen and Lisman, 1996; Sato and Yamaguchi, 2003; Tsodyks and Sejnowski, 1995) in which recurrent networks learn and strengthen forward sequences through forward movements, our model proposes goal-directed pathway learning through reverse sequences.

## RESULTS

### Hebbian plasticity with short-term depression potentiates reverse synaptic transmissions

We first simulated a sequential firing pattern that propagates through a one-dimensional recurrent neural network, and evaluated weight changes by Hebbian plasticity rules. The network consists of 500 rate neurons, which were connected with distance-dependent excitatory synaptic weights modulated by short-term synaptic plasticity (STP) (Romani and Tsodyks, 2015; Wang et al., 2014). In addition, the network had global inhibitory feedback to all neurons. A first external input to a neuron at one end (#0) elicited traveling waves of neural activity propagating to the opposite end (Figure 1A). Here we regard these activity patterns as a model of hippocampal firing sequences (Romani and Tsodyks, 2015; Wang et al., 2014). A second external input to a neuron at the center of the network triggered firing sequences propagating to both directions (Figure 1A) because synaptic weights were symmetric (Figure 1B). Here, we implemented a standard Hebbian plasticity rule, which potentiated excitatory synaptic weights by the product of postsynaptic and presynaptic neural activities. During the propagation of the first unidirectional sequence, this rule potentiated synaptic weights symmetrically without creating any bias in the synaptic weights (Figure 1B).

**Figure 1:**
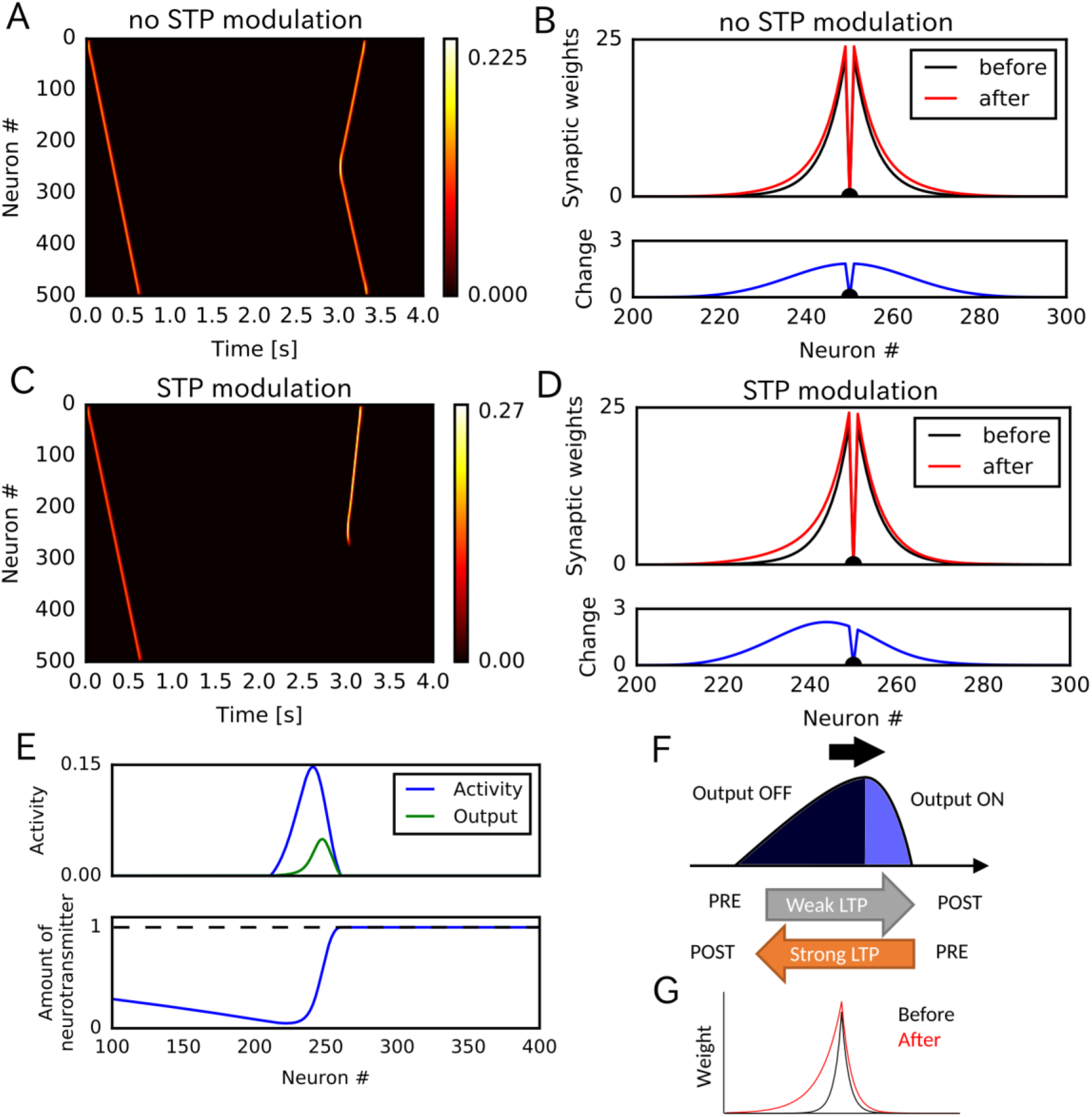
Potentiation of reverse propagation in 1-D recurrent network of rate neurons. (A, C) Weight modifications were induced by a firing sequence in a 1-D recurrent network (at time 0 sec), and the effects of these changes on sequence propagation were examined at time 3 sec. Recurrent synaptic weights were modified by the standard Hebbian plasticity rule (A) or the revised Hebbian plasticity rule, where the latter was modulated by STP. (B,D) Weights of outgoing synapses from neuron #250 to other neurons are shown at time 0 seconds (black) and 3 seconds (red) of the simulations in (A) and (C), respectively (top). Lower panels show the weight changes (blue). (E) Neural activity, presynaptic outputs (top) and the amount of neurotransmitters at presynaptic terminals (bottom) are shown at 300 ms of the simulation in (C). (F) Activity packet traveling on the 1-D recurrent network and the resultant weight changes by the revised Hebbian plasticity rule are schematically illustrated. (G) Distributions of synaptic weights are schematically shown before and after a sequence propagation.

However, when synaptic weights were changed by a modified Hebbian plasticity rule (Experimental Procedure), in which the long-term plasticity is also regulated by STP at presynaptic terminals (Froemke et al., 2006), the second firing sequence only propagated to the reverse direction of the first one (Figure 1C). This selective propagation occurred because the first firing sequence potentiated synaptic weights asymmetrically in the forward and reverse directions, thus creating a bias in the spatial distribution of synaptic weights (Figure 1D). This means that firing sequences strengthen the reverse synaptic transmissions more strongly than the forward ones in this model, and reverse sequences are more likely to be generated after forward sequences.

Why does this model generate such a bias to the reverse direction? To explain this, we show the packet of neural activity during the first firing sequence (at *t* = 300 ms) in Figure 1E, top. We also plotted neurotransmitter release from the presynaptic terminal of each neuron (presynaptic outputs), which is determined by the product of presynaptic neural activity and the amount of available neurotransmitters in the combined Hebbian rule (Figure 1E, bottom). Because neurotransmitters are exhausted in the tail of the activity packet, presynaptic outputs are effective only at the head of the activity packet. Due to this spatial asymmetricity of presynaptic outputs, connections from the head to the tail are strongly potentiated but those from the tail to the head are not (Figure 1F). This results in the biased potentiation of reverse synaptic transmissions in this model (Figure 1G). In other words, the packet of presynaptic outputs becomes somewhat “prospective” (namely, slightly skewed toward the direction of activity propagation), so the weight changes based on the coincidences between presynaptic outputs and postsynaptic activities result in the selective potentiation of connections from the “future” to the “past” in the firing sequence. This mechanism was not known previously and enables forward sequences to potentiate the synaptic transmissions responsible for reverse replay, and vice versa.

## Potentiation of reverse synaptic transmissions by STDP

The potentiation of reverse synaptic transmissions also occur robustly in spiking neurons with STDP. To show this, we constructed a one-dimensional recurrent network of Izhikevich neurons (Izhikevich, 2003; Izhikevich et al., 2004) connected via conductance-based AMPA and NMDA synaptic currents (see Methods). Initial synaptic weights, STP, and global inhibitory feedback were similar to those used in the rate neuron model. We tested two types of STDP: Hebbian STDP in which pre-to-post firing leads to potentiation and post-to-pre firing leads to depression, and symmetric STDP in which both firing orders result in potentiation if two spikes are temporally close or depression if two spikes are temporally distant. Experimental evidence suggests that recurrent synapses in hippocampal CA3 undergo symmetric STDP (Mishra et al., 2016). Contributions of all spike pairs were taken into account in these simulations.

Among these STDP types, only symmetric STDP showed a similar effect to the rate-based Hebbian rule. First, we simulated STDP without modulations by STP. The model propagated a spike sequence when a first stimulus is given to one end of the chain (Figure 2A). As expected, the firing sequence strengthened the forward sequence propagation (Figure 2A) and the related synaptic connections under Hebbian STDP (Figure 2B). Under symmetric STDP, synaptic transmissions were potentiated in both directions (Figure 2C and 2D). Introducing modulation of STDP by STP did not change the qualitative results under Hebbian STDP (Figure 2E and 2F). By contrast, a modified symmetric STDP modulated by STP biased the weight changes to the reverse direction (Figure 2H), and accordingly the second firing sequence selectively traveled towards the opposite direction to the first sequence (Figure 2G). These results indicate that a greater potentiation of reverse synaptic transmissions occurs in CA3 under the modified symmetric STDP.

**Figure 2:**
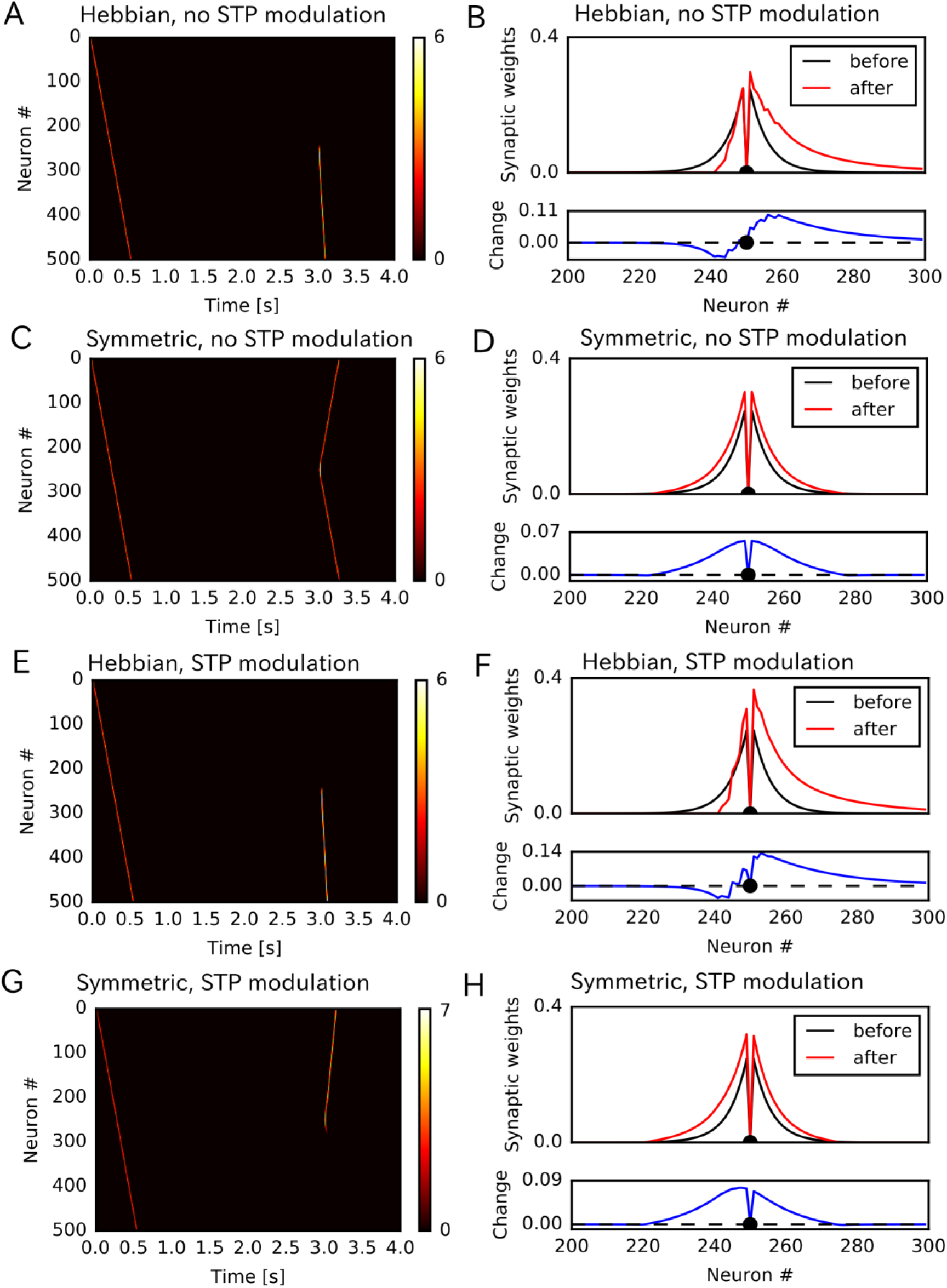
Potentiation of reverse propagation in 1-D recurrent network of spiking neurons. (A,C,E,G) Simulations similar to those shown in Figure 1 were performed in a 1-D spiking neural network. Recurrent synaptic weights were changed by Hebbian STDP (A,E) or symmetric STDP (C,G). In (A) and (B) the plasticity rules were not modulated by STP whereas in (C) and (D) the rules were modulated. (B,D,F,H) The weights of outgoing synapses from neuron #250 (top) are shown at time 0 seconds (black) and 3 seconds (red) of the simulation settings shown in (A), (C), (E) and (G), respectively. Lower panels display the weight changes (blue).

## Evaluation of parameter dependence in Poisson spike trains

We further confirmed the bias effect of the modified symmetric STDP in broader conditions than in the above network simulation. To this end, we generated sequential firing patterns along the one-dimensional network by sampling from a Poisson process, while manually controlling the number of propagating spikes per neuron, the mean inter-spike interval (ISI) of Poisson input spike trains, and time lags in spike propagation (i.e., time difference between the first spikes of neighboring neurons) (Figure 3A). The amount of neurotransmitter release by each presynaptic spike was calculated by the STP rule (Experimental Procedure), and the magnitude of long-term synaptic changes was calculated by a Gaussian-shaped symmetric STDP (Figure 3B) (Mishra et al., 2016). The net effect of synaptic plasticity was given as the product of the two quantities, as in the previous simulations. The parameter values of short-term and long-term plasticity were adopted from experimental results (Figure 3B and 3C) (Guzman et al., 2016; Mishra et al., 2016). We calculated the long-term weight changes in synapses sent from the central neuron in the network, and defined a weight bias as the difference in synaptic weights between forward and reverse directions (in which positive values mean bias to the reverse direction). We then obtained the mean bias and the fraction of positive biases (P(bias>0)) over 100 different realizations of spike trains generated with the same parameter values. For each number of spikes per neuron (2, 3, 4 and 5 spikes), we ran 1000 simulations using different mean ISIs and time lags randomly sampled from the interval [5 ms, 50 ms].

**Figure 3:**
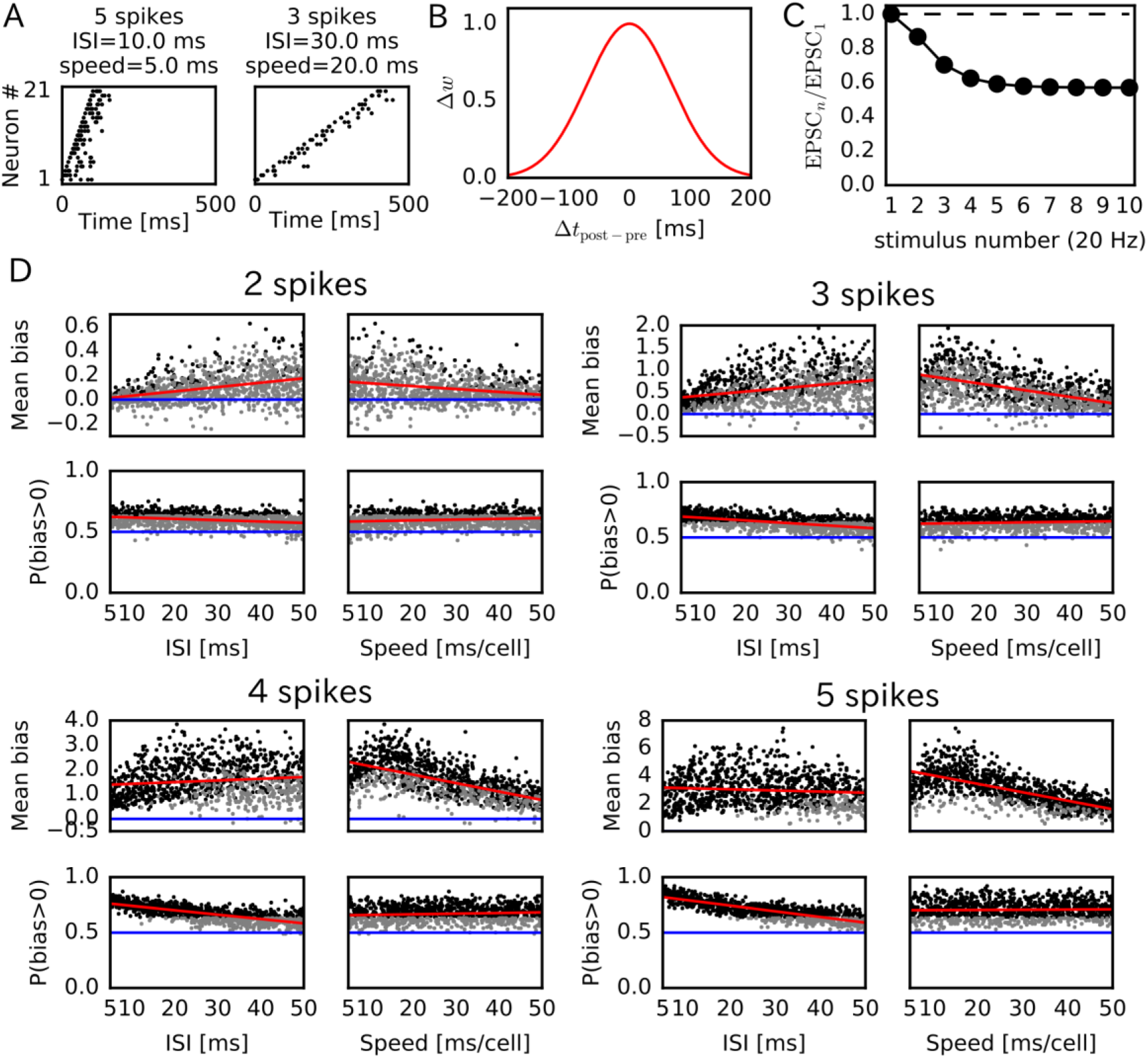
Reversely biased weight changes for symmetric STDP against variations in firing patterns. (A) Two examples show firing sequences having different ISIs and propagation speeds. (B) Gaussian-shaped symmetric STDP was used in the simulations (c.f., Figure 1 in Mishra et al., 2016). (C) Changes in the amplitude ratio of EPSC during a 20 Hz stimulation are shown (c.f., Figure S5 in Guzman et al., 2016). See the text for the parameter setting. (D) The mean (top) and fraction (bottom) of biases towards reverse direction are plotted for various parameter settings. A more positive bias indicates stronger weight changes in the reverse direction. Black dots correspond to parameter settings giving statistically significant positive or negative biases (P<0.01 in Wilcoxon signed rank test for the mean bias or binomial test for P(bias>0)) whereas grey dots are not significant. Red lines are linear fits to the data and blue lines indicate zero-bias (mean bias=0 and P(bias>0)=0.5).

Significant biases towards the reverse direction were observed in broad simulation conditions (Figure 3D). The biases were statistically significant (P<0.01 in Wilcoxon signed rank test for mean bias or binomial test for P(bias>0)) already in a part of conditions with 2 spikes per neuron, and the effect became prominent as the number of spikes increased. In general, the magnitude of synaptic changes is larger for a faster spike propagation (Figure 3D, Supplementary Table 1), which is reasonable because the potentiation of symmetric STDP becomes stronger as presynaptic firing and postsynaptic firing get closer in time. On the other hand, P(bias>0) was greater for a smaller mean ISI, and it did not depend on the propagation speed (Figure 3D, Supplementary Table 1). Especially, all simulations showed statistically significant biases to the reverse direction regardless of the propagation speed when the number of spikes per neuron is 4 or 5 and the mean ISI < 20 ms. This implies that the bias towards the reverse direction is the most prominent when the neural network propagates a sequence of bursts with intraburst ISIs less than 20 ms. Such bursting is actually observed in CA3 in vivo (Mizuseki et al., 2012) and simulations of a CA3 recurrent network model suggest that bursting enhances propagation of firing sequences (Omura et al., 2015).

Parameters that regulate STP also influence the bias towards the reverse direction. Actually, these parameters largely change in the hippocampus depending on experimental settings (Guzman et al., 2016). Therefore, we also performed simulations with randomly sampled values of the initial release probability of neurotransmitters (*U*) and the time constants of short-term depression and facilitation (τ_STD_ and τ_STF_). Here, the number of spikes per neuron was fixed to five, and the ISI and time lag were independently sampled from the interval [5 ms, 20 ms] in every trial. As shown in Figure 4, both mean bias and P(bias>0) clearly depend on the initial release probability. Prominent bias was observed for *U* > 0.3, and it gradually disappeared as *U* was decreased. The bias was weakly correlated with τ_STD_, but there was almost no correlation between the bias and τ_STF_. We show quantitative evaluations in Supplementary Table 1. In sum, the biases towards the reverse direction occurred robustly for a sufficiently high intraburst firing frequency and a sufficiently high release probability of neurotransmitters.

**Figure 4:**
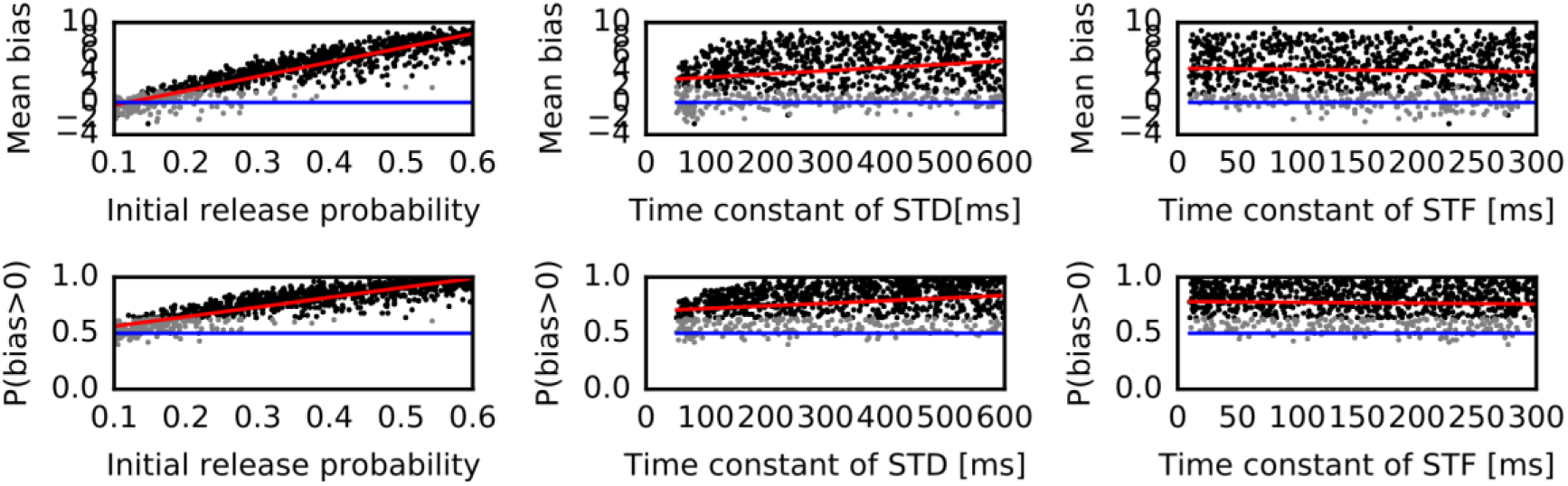
Reversely biased weight changes against variations in short-term plasticity. The mean and fraction of positive biases towards reverse direction are shown in various parameter settings of short-term plasticity, where a more positive bias indicates stronger weight changes in the reverse direction. Black dots correspond to parameter settings giving statistically significant positive or negative biases (P<0.01 in Wilcoxon signed rank test for the mean bias or binomial test for P(bias>0)) whereas grey dots are not significant. Red lines are linear fits to the data and blue lines indicate zero-bias (mean bias=0 and P(bias>0)=0.5).

In contrast to symmetric STDP, Hebbian STDP was not effective in potentiating reversed synaptic transmissions even if it was modulated by STP. In our simulations with parameters taken from experiments in CA1 (Bi and Poo, 2001), such a potentiation effect was never observed for Hebbian STDP with all-to-all spike coupling for any parameter value of STP and the number of spikes per neuron (5 or 15 spikes) (Supplementary Figure 1). However, Hebbian STDP with nearest-neighbor spike coupling (Izhikevich and Desai, 2003), in which only the nearest postsynaptic spikes before and after a presynaptic spike were taken into account, generated statistically significant biases towards the reverse direction in some parameter region (Supplementary Figure 2). In this case, large biases required large values of *U* and τ_STD_, and a large number of spikes per neuron (15 spikes). Thus, the condition that Hebbian STDP generates the biases to the reverse direction is severely limited although we cannot exclude this possibility.

## Goal-directed pathway learning through reverse replay

We now demonstrate how reverse replay events starting from a rewarded position enables the learning of goal-directed paths. We consider the case where an animal is exploring on a T-maze (Figure 5A). During navigation, the animal gets a reward at the one end of the arm (position D2), but not at the opposite end (position D1) and other locations. In each trial, the animal starts at the center arm (position A) and runs into one of the two side arms at position B. In the present simulations, the animal visits both ends alternately: it reaches to D1 in (2*n* + 1)-th trials and D2 in (2*n*)-th trials, where *n* is an integer. After reaching either of the ends (i.e., D1 or D2), the animal stops there for 7 s.

**Figure 5:**
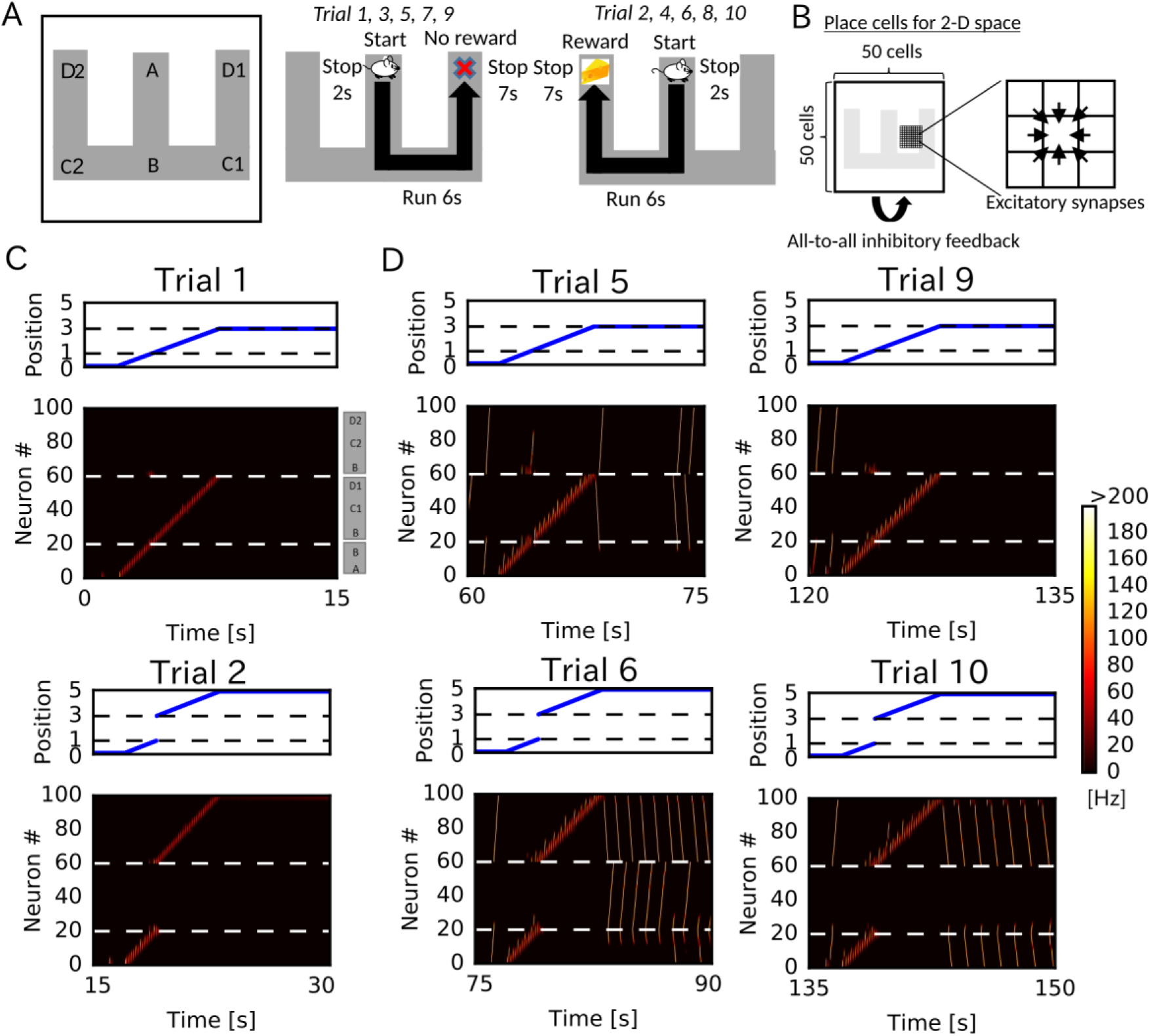
Goal-directed pathway learning with reverse replay on a T-maze. (A) An animal alternately repeats traverses from starting position A and two goal positions D1 and D2 on a track. (B) Setting of a 2-D recurrent network of place cells. The combined receptive fields of all place cells cover the entire track. Adjacent place cells were interconnected with excitatory synapses. (C) The position of the animal (top) and simulated neural activities of place cells on the track (bottom) in each trial. The positions on the T-shape track correspond to the ordinate as follows: A → B:0 to 1; B → D1:1 to 3; B → D2:3 to 5.

We constructed the 50 x 50 two-dimensional (2-D) place-cell network associated to the 2-D space that the animal explored (Figure 5B), using the rate neuron model. Each place cell had a place field in the corresponding position on the 2-D space and received global inhibitory feedback proportional to the overall network activity. Neighboring place cells were reciprocally connected with excitatory synapses, which were modulated by short-term and long-term plasticity rules as in Figure 1. During the delivery of reward, we mimicked dopaminergic modulations by enhancing the inputs to CA3 (Kobayashi and Suzuki, 2007) and increasing the frequency of triggering firing sequences (Ambrose et al., 2016; Singer and Frank, 2009). Under this condition, a larger number of reverse replay was generated in the rewarded position. Thus, larger potentiation of synaptic pathways towards reward is expected in our model.

In early trials, recurrent synaptic connections were relatively weak and not organized for the spatial navigation task. Therefore, the network only showed theta-modulated place-cell activity driven by external inputs the recurrent network during locomotion, and reverse replay and forward replay (prospective firing sequences) were rare (Figure 5C and Supplementary Movie 1). However, in later trials, the network generated reverse replay (at D1 and D2) and prospective firing sequences (at A) during immobility (Figure 5D and Supplementary Movie 1). Notably, in these trials, prospective firing sequences traveled only towards D2, where the animal had got reward. Furthermore, backward firing sequences that started from the non-rewarded position D1 propagated to the rewarded position D2 but not to the start (reverse replay). This implies that the network model forms synaptic pathways to propagate neural activity that guides the animal towards reward through a combination of paths that has not been traversed by the animal (i.e., the animal never traveled directly from D2 to D1 in our simulations). All these properties of firing sequences look convenient for the goal-directed learning of spatial map.

To visualize how the recurrent network was optimized for the goal-directed exploratory behavior, we defined “connection vectors” from recurrent synaptic weights. For each place cell, we calculated the weighted sum of eight unit vectors each directed towards one of the eight neighboring neurons, using the corresponding synaptic weights (Figure 6A). These connection vectors represent the average direction of neural activity transmitted from each neuron, and the 2-D vector field shows the flow of neural activities in the 2-D recurrent network and hence in the 2-D maze. We note that these vectors bias the flow, but actual firing sequences can sometimes travel in different directions from the vector flow. Initially, synaptic connections were random and the connection vector field was not spatially organized (Supplementary Figure 3). However, after the exploration, the vector field was organized so as to route neural activities to those neurons encoding the rewarded position on the track (Figure 6B). A similar route map was also obtained when we reversed the sequential order of visits to the two arms (Supplementary Figure 4), but was abolished when we removed reward (Supplementary Figure 5) or the effect of STP on Hebbian plasticity (Supplementary Figure 6). As demonstrated previously, direct synaptic pathways from D1 to D2 were also created. The emergence of direct paths relies on two mechanisms in this model. First, as seen in Supplementary Figure 5, connections are biased from goal to start when there is no reward at the goal because theta sequences enhance synaptic pathways opposite to the direction of animal's movement. In non-rewarded travels, these directional biases are not overwritten by reverse replay. Thus, the relative preference of a synaptic pathway in hippocampal sequential firing decreases for exploration that does not result in reward. Second, some of reverse replay sequences from D2 propagates into D1 instead of the stem arm (Figure 5D and Supplementary Movie 1), and such “joint replay” enhances biases towards goal through unexperienced pathways. Thus, our model creates a map not only for the spatial paths experienced by the animal, but also for their possible combinations if they guide the animal directly to the rewarded position from a point in the space. In this sense, our model optimizes the cognitive map of the spatial environment.

**Figure 6:**
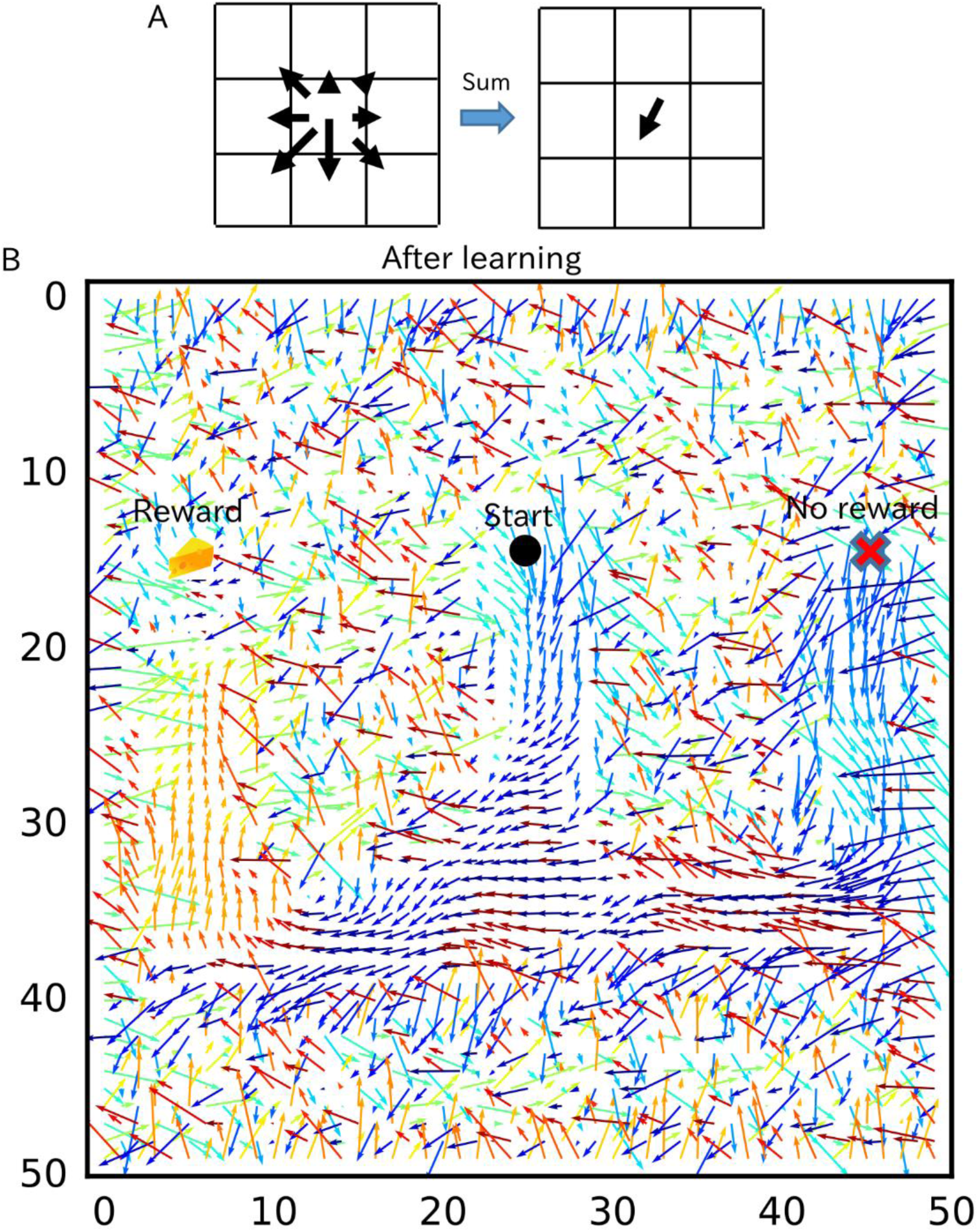
Recurrent connections organized on the T-maze for activity propagation towards reward. (A) Calculation of connection vectors at each position on the T-maze is schematically illustrated. Vector length in the left figure corresponds to synaptic weights. (B) The connection vectors formed through reverse replay point the directions leading to the rewarded goal along the track.

## DISCUSSION

In this paper, we showed that the modified Hebbian plasticity rule modulated by STP biases synaptic weights towards the reverse direction of the firing sequences that traveled through a recurrent network. We demonstrated this counterintuitive phenomenon in network models of rate neurons and those of spiking neurons obeying symmetric STDP. We further clarified for various Poisson-like sequential firing patterns that the phenomenon favors spike bursts and a high presynaptic release probability. We also showed that the selective potentiation of reverse directions is unlikely to occur for the conventional Hebbian STDP.

Our results have several implications for spatial memory processing by the hippocampus. Suppose that the animal is rewarded at a spatial position after exploring a particular path. Reverse replay propagating backward from the rewarded location will strengthen the neuronal wiring in CA3 that preferentially propagates forward firing sequences to this location along the path. Because the frequency of reverse replay increases at rewarded positions (Ambrose et al., 2016; Singer and Frank, 2009), reward delivery induces the preferential potentiation of forward pathways, which in turn results in an enhanced occurrence of forward replay in the consolidation phase. Thus, our model predicts that reverse reply is crucial for the reinforcement of reward-seeking behavior in the animal and gives, for the first time, the mechanistic account for the way reverse replay enables the hippocampal prospective coding of reward-seeking navigation. Our computational results are qualitatively consistent with the experimentally observed properties of forward and reverse replay events (Ambrose et al., 2016; Carr et al., 2011; Diba and Buzsáki, 2007; Foster and Wilson, 2006; Pfeiffer and Foster, 2013; Singer and Frank, 2009) and matches to the recent finding of symmetric STDP time windows in CA3 (Mishra et al., 2016).

The most critical assumption in our model is the rapid modulation of STDP coherent to the presynaptic neurotransmitter release of STP. Such a modulation was actually reported in the visual cortex (Froemke et al., 2006). Although short-term depression also exists in CA3 (Guzman et al., 2016), STDP is modulated in a slightly different fashion in the hippocampus: the strong modulation arises from the second presynaptic spike rather than the first one (Wang et al., 2005). However, the experiment was performed in a dissociated culture in which hippocampal sub-regions were not distinguished. Moreover, the modulation of STDP was not tested for more than two presynaptic spikes, and whether a third presynaptic spike further facilitates or rather depresses STDP remains unknown. Therefore, the contributions of STP to STDP should be further validated in CA3. The proposed role of short-term depression in biasing replay events may also be examined by pharmacological blockade or enhancement of STP in the hippocampus (Froemke et al., 2006).

Neuromodulations, especially reward-triggered facilitation of replay events (Ambrose et al., 2016; Singer and Frank, 2009), play important roles for the proposed mechanism of goal-directed learning with reverse replay. Dopamine enhances neuronal excitability and synaptic plasticity in CA1 and dentate gyrus (Lisman and Grace, 2005; Lisman et al., 2011; Yang and Dani, 2014) and optogenetic stimulation of dopaminergic fibers from VTA during exploration promotes subsequent reactivation of CA1 cell assemblies for memory persistence (McNamara et al., 2014). On the other hand, CA3 primarily receives dopaminergic input from the locus coeruleus (Walling et al., 2012), which signals novelty to hippocampus (Takeuchi et al., 2016). Therefore, learning in CA3 may be affected by not reward but other information such as novelty and salience of sensory inputs (Lisman and Grace, 2005; Lisman et al., 2011). However, even if reward-induced dopamine is not effective in CA3 recurrent synapses, dopamine enhances inputs from dentate gyrus to CA3 (Kobayashi and Suzuki, 2007), which may increase the frequency of reverse replay and suffice for the goal-directed learning. In addition, inputs from dentate gyrus may provide a reinforcement signal for potentiation of CA3 recurrent synapses (Kobayashi and Poo, 2004). Our model also predicts that the release probability of neurotransmitter strongly affects the magnitude and probability of bias towards the reverse direction (Figure 4). Therefore, modulations of neurotransmitter release in CA3 can regulate the behavioral impact of hippocampal firing sequences. For example, acetylcholine suppresses neurotransmitter release at recurrent synapses (Hasselmo, 2006), which may abolish the directional biases created by theta sequences (see Figure 4 and Supplementary Figure 5). Presynaptic long-term plasticity (Costa et al., 2015) also affects the directional biases at longer timescales.

Importantly, our model reinforces unexperienced paths by connecting the multiple paths that were previously reinforced by separate experiences. In the simulations of the T-maze task, the network model not only learned actually traversed paths from a start (A) to a goal (D2), but also remembered paths from other locations (C1 and D1) to the goal despite that the animal had not experienced these paths. This reinforcement occurs because reverse replay sequences starting from the visited arm occasionally propagate or bifurcate into an unvisited arm at the branching point. In the hippocampus, two replay sequences were jointly observed in the forward and reverse directions along the two arms of a Y-shape track (Wu and Foster, 2014), suggesting that the above sequence propagations are possible. If, however, reverse replay, hence pathway learning, only occur on recently experienced paths, we may transiently upregulate the excitability of most recently activated neurons, as in the previous models of reverse replay (Foster and Wilson, 2006; Molter et al., 2006).

While the present model could demonstrate goal-directed pathway learning on a T-maze, the model has to be yet improved to learn more complex tasks such as navigation on an alternating T-maze. In the present T-maze task, we simulated only the relatively easy cases in which reward was always delivered at the same location. However, in an alternating T-maze task, the animal has to remember recent experiences to change its choices based on the stored memory. Many possible extensions of the model will achieve this task. A straightforward extension is to maintain multiple charts representing the alternating paths (Samsonovich and McNaughton, 1997) and switch them according to the stored short-term memory or certain context information. Each chart will be selectively reinforced by reverse replay along one of the alternating paths.

Reverse replay has not been found in the neocortical circuits. For instance, firing sequences in the rodent prefrontal cortex are reactivated only in the forward directions (Euston et al., 2007). To the best of our knowledge, neocortical synapses obey Hebbian STDP (with asymmetric time windows) (Froemke et al., 2006). This seems to be consistent with the selective occurrence of forward sequences because, as suggested by our model, the sensory-evoked forward firing sequences should only strengthen forward synaptic pathways under Hebbian STDP. However, if dopamine turns Hebbian STDP into symmetric STDP, which is actually the case in hippocampal area CA1 (Brzosko et al., 2015; Zhang et al., 2009), forward firing sequences will reinforce the propagation of reverse sequences. Whether reverse sequences exist in the neocortex and, if not, what functional roles replay events play in the neocortical circuits are still open questions.

Whether the present neural mechanism to combine experienced paths into a novel path accounts for cognitive functions other than memory is an intriguing question. For instance, does this mechanism explain the transitivity rule of inference by neural networks? The transitive rule is one of the fundamental rules in logical thinking and says, “if A implies B and B implies C, then A implies C.” This flow of logic has some similarity to that of joint forward-replay sequences, which says, “if visiting A leads to visiting B and visiting B leads to visiting C, then visiting A leads to visiting C.” Logic thinking is more complex and should be more rigorous than spatial navigation, and little is known about its neural mechanisms. The proposed neural mechanism of pathway learning may give a cue for exploring the neural substrate for logic operations by the brain.

In sum, our model proposes a biologically plausible mechanism for the global search of optimal forward paths through the rapid modulation of long-term plasticity by short-term plasticity effects. In the dynamic programming-based path finding methods such as Dijkstra’s algorithm for finding the shortest path (Dijkstra, 1959) and Viterbi algorithm for finding the most likely state sequences in a hidden Markov model (Bishop, 2010), a globally optimal path is determined by backtrack from the goal to the start after an exhaustive local search of forward paths. Our model enables similar path finding mechanism through reverse replay. Such a mechanism has been suggested in machine learning literature (Foster and Knierim, 2012), but whether and how neural dynamics achieves it remained unknown. In addition, our model predicts the roles of neuromodulators that modify plasticity rules and activity levels in sequence learning. These predictions are testable by physiological experiments.

## METHODS

### One-dimensional recurrent network model with rate neurons (Figure 1)

We simulated the network of 500 neurons. Firing rate of neuron *i* were determined as

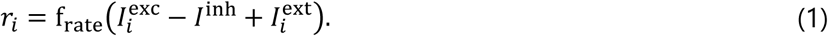

The function f_rate_(*I*) was threshold linear function

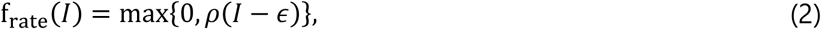

where *ρ* = 0.0025 and *ε* = 0.5. Excitatory synaptic current 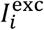 and Inhibitory feedback 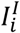 followed

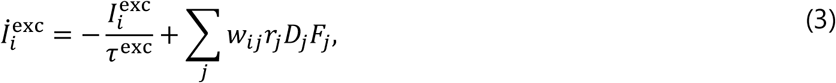

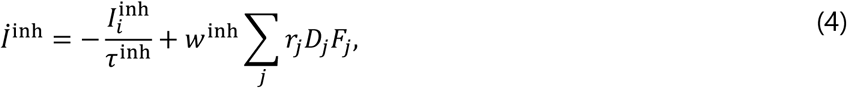

where *τ*^exc^ = *τ*^inh^ = 10 ms, and *w*^inh^ = 1. Variables for short-term synaptic plasticity *D*_*j*_ and *F*_*j*_ were calculated as (Wang et al., 2014)

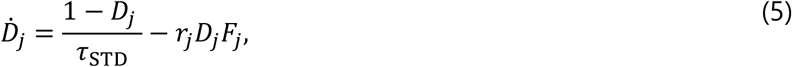

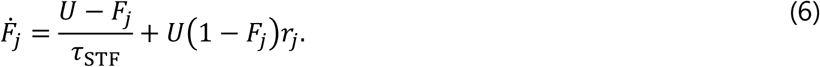

Parameters were *τ*_STD_ = 500 ms,*τ*_STF_ = 200 ms, and *U* = 0.6. External input 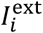 was usually zero and changed to 5 for 10 ms when the cell was stimulated. We stimulated neurons 0 ≤ *i* ≤ 10 at the beginning of simulations, and neurons 245 ≤ *i* ≤ 255 at 3 seconds after the beginning.

Initial values of excitatory weights *w*_ij_ were determined as

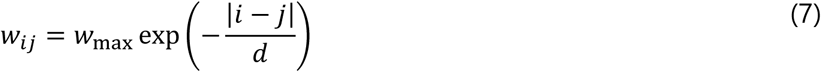

where *w*_max_ was 27 and *d* = 5. Self-connections w_ii_ were set to zero throughout the simulations.

The weights were modified according to Hebbian synaptic plasticity as

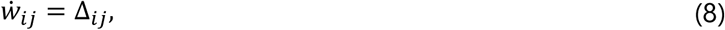

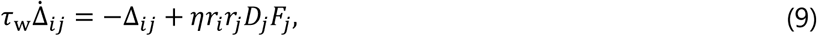

where *η* = 20 and *τ*_w_ = 1000 ms. When we simulated Hebbian synaptic plasticity without the modulation by short-term plasticity, *D*_*j*_*F*_*j*_ was removed from this equation and *η* value was changed to 4.

### One-dimensional recurrent network model with spiking neurons (Figure 2)

We used Izhikevich model (Izhikevich, 2003) for these simulations:

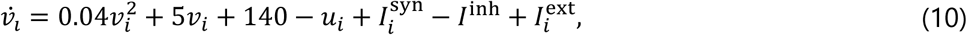

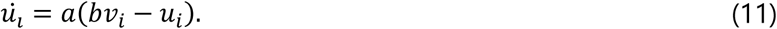

If the membrane voltage *v*_*i*_ ≥ 30 mV, the neuron emits a spike and the two variables were reset as *v*_i_ ← *c* and *u*_*i*_ ← *u*_*i*_ + *d*. Parameter values were *a* = 0.02, *b* = 0.2, *c* = -65 and *d* = 8. Excitatory synaptic current was

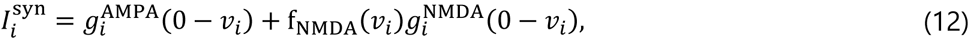

and the synaptic conductances followed

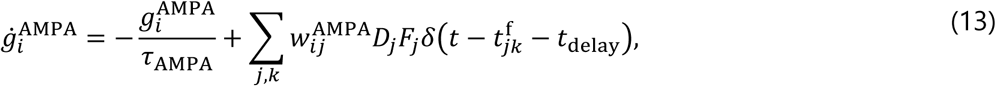

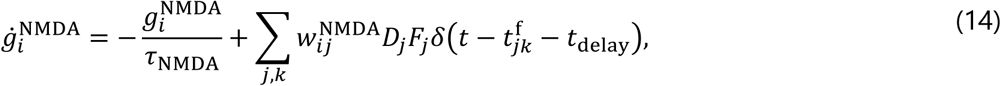

where 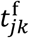 is the timing of the *k*-th spike of neuron *j* and parameter values were set as *τ*_AMPA_ = 5 ms, *τ*_NMDA_ = 150 ms and *t*_delay_ = 2 ms. The voltage dependence of NMDA current (Izhikevich et al., 2004) was described as

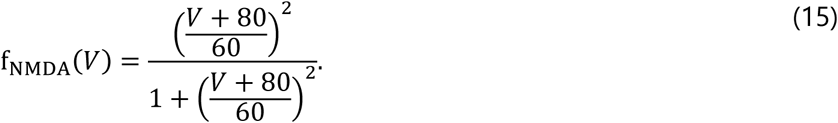

Inhibitory feedback 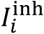 was given as

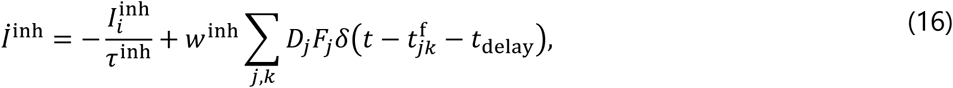

where *w*^inh^ = 1 and *τ*^inh^ = 10 ms. Short-term synaptic plasticity was described as (Wang et al., 2014)

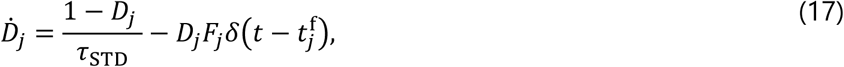

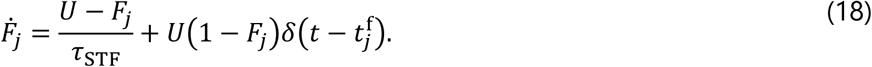

Parameter values were *τ*_STD_ = 500 ms,*τ*_STF_ = 200 ms and *U* = 0.6. External input 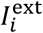 was the same as that in the rate neuron model.

Initial values of synaptic weights 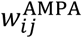 were determined as

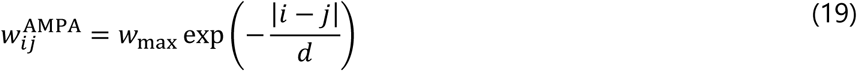

where *w*_max_ was 0.3 and *d* = 5. Self-connections *w*_*ii*_ were set to zero throughout the simulations. The weights of NMDA current 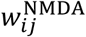 were determined as 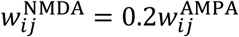 and fixed at these values throughout the simulations.

The weights of AMPA current were modified by STDP as

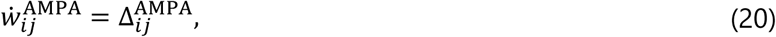

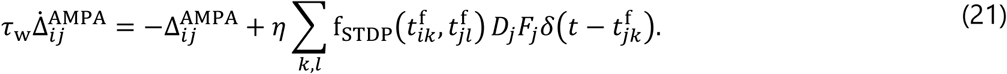

We simulated two different STDP types by changing the function f_STDP_(*t*_post_, *t*_pre_) as follows.

Hebbian STDP:

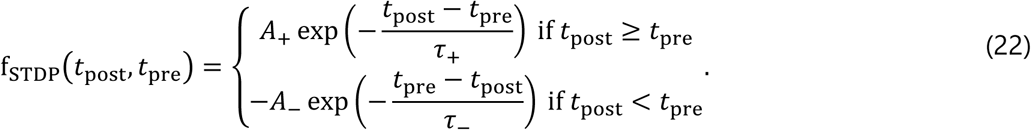

Symmetric STDP:

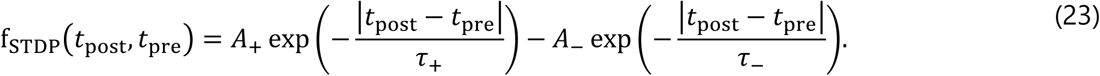

Parameter values were set as *η* = 0.05, *τ*_w_ = 1000 ms, *A*_+_ = 1, *A*_-_ = 0.5, *τ*_+_ = 20 ms and *τ*_-_ = 40 ms for all STDP types. When we simulated STDP without the modulation by short-term plasticity, *D*_*j*_*F*_*j*_ was removed from the above equation and the *η* value was changed to 0.01.

### Performance evaluation with poisson spike trains (Figures 3 and 4)

In each simulation, we sampled spike trains of 21 neurons (neuron #1 - #21) 100 times for given values of the number of spikes per neuron N_spike_, mean inter-spike interval (ISI) *t*_ISI_, and firing propagation speed *t*_speed_. First spikes of neuron #n was set to 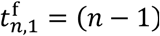*t*_speed_. Following a first spike, Poisson spike train for each neuron was simulated by sampling ISI 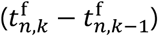 from the following exponential distribution:

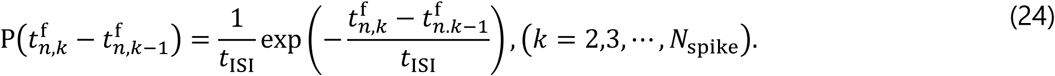

We induced an absolute refractory period by resampling ISI if it was shorter than 1 ms. After we obtained spike trains, neurotransmitter release was simulated by solving Equations (17) and (18) for each neuron. In Figure 3, parameter values of short-term plasticity were set as *τ*_STD_ = 150 ms,*τ*_STF_ = 40 ms and U = 0.37. In Figure 4, we evaluated biases for the values of *U*,*τ*_STD_ and *τ*_STF_ sampled from [0.1, 0.6], [50 ms, 500 ms] and [10ms, 300 ms], respectively.

Changes in the weight from neuron #11 in the center (*j* = 11) to neuron i were calculated as

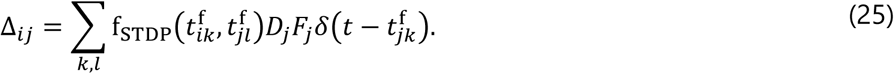

In all-to-all STDP, we calculated the above summation over all spike pairs. In nearest-neighbor STDP (Izhikevich and Desai, 2003), we considered only pairs of a presynaptic spike and the nearest postsynaptic spikes before and after the presynaptic spike. For symmetric STDP, f_STDP_(*t*_post_, *t*_pre_) was Gaussian (Mishra et al., 2016)

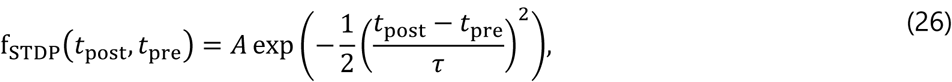

where *A* = 1 and *τ* = 70 ms. For Hebbian STDP, f_STDP_(*t*_post_, *t*_pre_) was the same as the previous one except that parameter values were changed as *A*_+_ = 0.777,*A_-_* = 0.273, *τ*_+_ = 16.8 ms and *τ_-_* = 33.7 ms (Bi and Poo, 2001). Weight biases were calculated for each spike train as 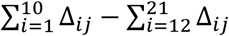.

### Simulation of the goal-directed learning in T-maze (Figures 5 and 6)

We simulated an animal moving on a two-dimensional space spanned by *x* and (0 ≤ *x* ≤ 50,0 ≤ *y* ≤ 50). Coordinates of the six corners (A, B, C1, C2, D1, D2) of T-maze were **z**_A_ = (25, 15), **z**_B_ = (25,35), **z**_C1_ = (45,35), **z**_C2_ = (45,15), **z**_D1_ = (5,35) and **z**_D2_ = (5,15). In each set of trials with time length *T* = 15s, the position of the animal **z**_pos_ at *t*′ = *t* mod *T* was determined as

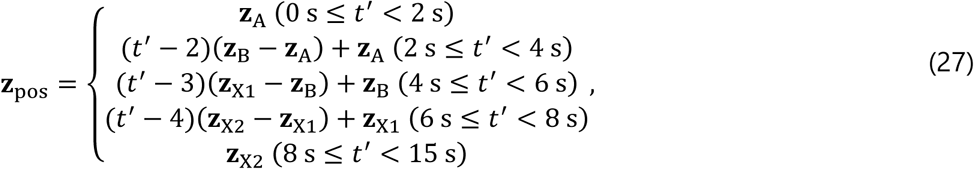

where **z**_X1_ = **z**_C1_ and **z**_X2_ = **z**_C2_ in the (2*n*+1)-th trials and **z**_X1_ = **z**_D1_ and **z**_X2_ = **z**_D2_ in the (2*n*)-th trials.

The neural network consisted of 2500 place cells that were arranged on a 50 x 50 two-dimensional square lattice. The place field centers of neurons in the *i*-th column (x-axis) and the *j*-th row (y-axis) were denoted as **Z**_*i,j*_ = (*i,j*). Each place cell received excitatory connections from eight surrounding neurons. The connection weight from a neuron at (*i-k,j-l*) to a neuron at (*i,j*) were denoted as 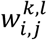, where the possible combinations of (*k, l*) were given as *S* = {(1,1), (1,0), (1,-1), (0,1), (0,-1), (-1,1), (-1,0), (-1,-1)}. Initial connection weights were uniformly random and normalized such that the sum of eight connections obeys 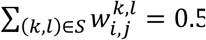

Activities of place cells *r*_*i,j*_ were simulated as

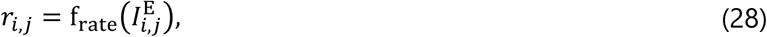

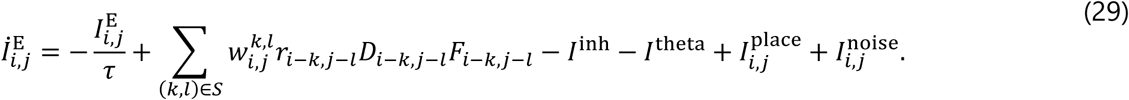

Time constant *τ* was 10 ms. The function f_rate_(*I*) was a threshold linear function

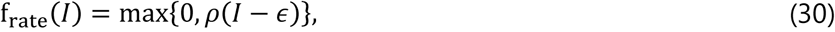

where *ρ* = 1 and *ε* = 0.002. Inhibitory feedback *I*^inh^ was simulated as

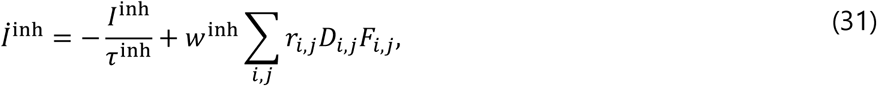

where *τ*^inh^ = 10 ms and *w*^inh^ = 0.0005. Variables for short-term synaptic plasticity *D*_*i,j*_ and *F*_*i,j*_ obeyed

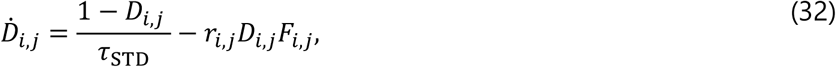

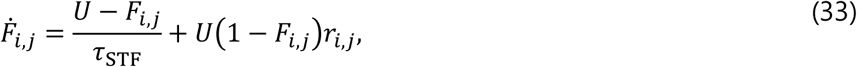

with parameter values *τ*_STD_ = 300 ms, *τ*_STF_ = 200 ms, and U = 0.4. Theta oscillation was induced by

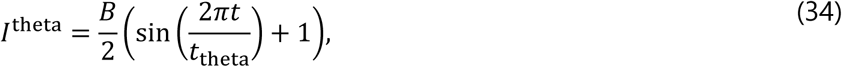

where B = 0.005 kHz and 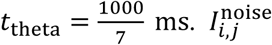 was independent Gaussian noise with the standard deviation 0.0005 kHz. Place-dependent inputs for each neuron 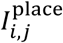 were determined from the place field center of each neuron **Z**_*i,j*_ and the current position of the animal **z**_pos_ as

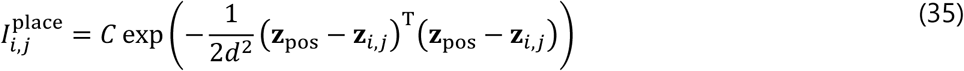

where *d* = 2. The parameter *C* was set as 0.005 kHz when the animal was moving and 0.001 kHz when the animal was stopping at D2 (the position of reward). When the animal was stopping at other positions, C was set at zero but occasionally changed to 0.001 kHz for a short interval of 200ms. The occurrence of this brief activation followed Poisson process at 0.1 Hz, but it always occurred one second after the onset of each trial to trigger prospective firing sequences.

Hebbian synaptic plasticity was implemented as

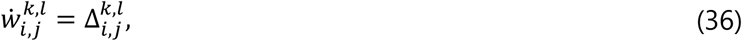

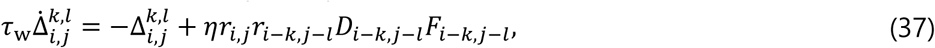

where *η* = 1 and *τ*_w_ = 30 s. If the sum of synaptic weights on each neuron 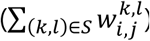 was greater than unity, we renormalized synaptic weights by dividing them by the sum. When we simulated Hebbian synaptic plasticity without modulations by short-term plasticity, *D*_*i-k,j-l*_*F*_*i-k,j-l*_ was removed from the above equation and the η value was changed to 0.1.

“Connection vector” of each neuron **u**_*i,j*_ was calculated as the following weighted sum of unit vectors 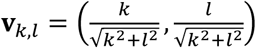

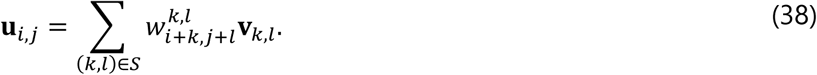

## CODE AVAILABLITY

Simulations and visualization were written in C++ and Python 3, and the codes are available at https://github.com/TatsuyaHaga/reversereplaymodel_codes.

## AUTHOR CONTRIBUTIONS

T. H. and T. F. equally contribute to the conception and design of the model as well as the preparation of the manuscript. T. H. performed numerical simulations and data analyses.

## ACKNOWLEDGEMENTS

We are grateful to Masami Tatsuno for fruitful discussion. This work was partly supported by Grants-in-Aid for Scientific Research (KAKENHI) from MEXT (15H04265, 16H01289, 17H06036) and CREST, JST (JPMJCR13W1).

## COMPETING INTERESTS STATEMENT

Authors declare no competing interests.

## REFRENCES

Ambrose, R.E., Pfeiffer, B.E., and Foster, D.J. (2016). Reverse Replay of Hippocampal Place Cells Is Uniquely Modulated by Changing Reward. Neuron 91, 1124–1136.

Bi, G., and Poo, M. (2001). Synaptic modification by correlated activity: Hebb’s postulate revisited. Annu. Rev. Neurosci. 24, 139–166.

Bishop, C.M. (2010). Pattern Recognition and Machine Learning (Springer).

Blum, K.I., and Abbott, L.F. (1996). A Model of Spatial Map Formation in the Hippocampus of the Rat. Neural Comput. 8, 85–93.

Brzosko, Z., Schultz, W., and Paulsen, O. (2015). Retroactive modulation of spike timingdependent plasticity by dopamine. Elife 4, 1–13.

Carr, M.F., Jadhav, S.P., and Frank, L.M. (2011). Hippocampal replay in the awake state: a potential substrate for memory consolidation and retrieval. Nat. Neurosci. 14, 147–153.

Costa, R.P., Froemke, R.C., Sjöström, P.J., and van Rossum, M.C.W. (2015). Unified pre- and postsynaptic long-term plasticity enables reliable and flexible learning. Elife 4, 1–16.

Diba, K., and Buzsáki, G. (2007). Forward and reverse hippocampal place-cell sequences during ripples. Nat. Neurosci. 10, 1241–1242.

Dijkstra, E.W. (1959). A Note on Two Problems in Connexion with Graphs. Numer. Math. 271, 269–271.

Euston, D.R., Tatsuno, M., and McNaughton, B.L. (2007). Fast-Forward Playback of Recent Memory Sequences in Prefrontal Cortex During Sleep. Science 318, 1147–1150.

Foster, D.J., and Knierim, J.J. (2012). Sequence learning and the role of the hippocampus in rodent navigation. Curr. Opin. Neurobiol. 22, 294–300.

Foster, D.J., and Wilson, M. a (2006). Reverse replay of behavioural sequences in hippocampal place cells during the awake state. Nature 440, 680–683.

Froemke, R.C., Froemke, R.C., Tsay, I.A., Raad, M., Long, J.D., and Dan, Y. (2006). Contribution of Individual Spikes in Burst-Induced Long-Term Synaptic Modification. J. Neurophysiol. 95, 1620–1629.

Gerstner, W., and Abbott, L.F. (1997). Learning navigational maps through potentiation and modulation of hippocampal place cells. J. Comput. Neurosci. 4, 79–94.

Guzman, S.J., Schlögl, A., Frotscher, M., Jonas, P., Eichenbaum, H., Kesner, R.P., McNaughton, B.L., Morris, R.G.M., Treves, A., Rolls, E.T., et al. (2016). Synaptic mechanisms of pattern completion in the hippocampal CA3 network. Science 353, 109–120.

Hasselmo, M.E. (2006). The role of acetylcholine in learning and memory. Curr. Opin. Neurobiol. 16, 710–715.

Izhikevich, E.M. (2003). Simple model of spiking neurons. IEEE Trans. Neural Networks 14, 1569–1572.

Izhikevich, E.M. (2007). Solving the distal reward problem through linkage of STDP and dopamine signaling. Cereb. Cortex 17, 2443–2452.

Izhikevich, E.M., and Desai, N.S. (2003). Relating STDP to BCM. Neural Comput. 15, 1511–1523.

Izhikevich, E.M., Gally, J.A., and Edelman, G.M. (2004). Spike-timing dynamics of neuronal groups. Cereb. Cortex 14, 933–944.

Jahnke, S., Timme, M., and Memmesheimer, R.-M. (2015). A Unified Dynamic Model for Learning, Replay, and Sharp-Wave/Ripples.J. Neurosci. 35, 16236–16258.

Jensen, O., and Lisman, J.E. (1996). Hippocampal CA3 region predicts memory sequences: accounting for the phase precession of place cells. Learn. Mem. 3, 279–287.

Kobayashi, K., and Poo, M.M. (2004). Spike Train Timing-Dependent Associative Modification of Hippocampal CA3 Recurrent Synapses by Mossy Fibers. Neuron 41, 445–454.

Kobayashi, K., and Suzuki, H. (2007). Dopamine selectively potentiates hippocampal mossy fiber to CA3 synaptic transmission. Neuropharmacology 52, 552–561.

Lee, A.K., and Wilson, M.A. (2002). Memory of sequential experience in the hippocampus during slow wave sleep. Neuron 36, 1183–1194.

Lisman, J.E., and Grace, A.A. (2005). The hippocampal-VTA loop: Controlling the entry of information into long-term memory. Neuron 46, 703–713.

Lisman, J., Grace, A.A., and Duzel, E. (2011). A neoHebbian framework for episodic memory; role of dopamine-dependent late LTP. Trends Neurosci. 34, 536–547.

McNamara, C.G., Tejero-Cantero, Á., Trouche, S., Campo-Urriza, N., and Dupret, D. (2014). Dopaminergic neurons promote hippocampal reactivation and spatial memory persistence. Nat. Neurosci. 17, 1658–1660.

Mishra, R.K., Kim, S., Guzman, S.J., and Jonas, P. (2016). Symmetric spike timing-dependent plasticity at CA3-CA3 synapses optimizes storage and recall in autoassociative networks. Nat. Commun. 7, 11552.

Mizuseki, K., Royer, S., Diba, K., and Buzsáki, G. (2012). Activity dynamics and behavioral correlates of CA3 and CA1 hippocampal pyramidal neurons. Hippocampus 22, 1659–1680.

Molter, C., Sato, N., Salihoglu, U., and Yamaguchi, Y. (2006). How reward can induce reverse replay of behavioral sequences in the hippocampus. Int. Conf. Neural Inf. Process. 1–10.

Nakashiba, T., Buhl, D.L., McHugh, T.J., and Tonegawa, S. (2009). Hippocampal CA3 Output Is Crucial for Ripple-Associated Reactivation and Consolidation of Memory. Neuron 62, 781–787.

Omura, Y., Carvalho, M.M., Inokuchi, K., and Fukai, T. (2015). A Lognormal Recurrent Network Model for Burst Generation during Hippocampal Sharp Waves. J. Neurosci. 35, 14585–14601.

Pfeiffer, B.E., and Foster, D.J. (2013). Hippocampal place-cell sequences depict future paths to remembered goals. Nature 497, 74–79.

Romani, S., and Tsodyks, M. (2015). Short-term plasticity based network model of place cells dynamics. Hippocampus 25, 94–105.

Samsonovich, A., and McNaughton, B.L. (1997). Path integration and cognitive mapping in a continuous attractor neural network model. J. Neurosci. 17, 5900–5920.

Sato, N., and Yamaguchi, Y. (2003). Memory Encoding by Theta Phase Precession in the Hippocampal Network. Neural Comput. 15, 2379–2397.

Singer, A.C., and Frank, L.M. (2009). Rewarded Outcomes Enhance Reactivation of Experience in the Hippocampus. Neuron 64, 910–921.

Takeuchi, T., Duszkiewicz, A.J., Sonneborn, A., Spooner, P.A., Yamasaki, M., Watanabe, M., Smith, C.C., Fernández, G., Deisseroth, K., Greene, R.W., et al. (2016). Locus coeruleus and dopaminergic consolidation of everyday memory. Nature 537, 357–362.

Tsodyks, M., and Sejnowski, T. (1995). Associative Memory and Hippocampal Place Cells. Int. J. Neural Syst. 6, 81–86.

Walling, S.G., Brown, R.A., Miyasaka, N., Yoshihara, Y., and Harley, C.W. (2012). Selective wheat germ agglutinin (WGA) uptake in the hippocampus from the locus coeruleus of dopamine-β-hydroxylase-WGA transgenic mice. Front. Behav. Neurosci. 6, 1–8.

Wang, H.-X., Gerkin, R.C., Nauen, D.W., and Bi, G.-Q. (2005). Coactivation and timing-dependent integration of synaptic potentiation and depression. Nat. Neurosci. 8, 187–193.

Wang, Y., Romani, S., Lustig, B., Leonardo, A., and Pastalkova, E. (2014). Theta sequences are essential for internally generated hippocampal firing fields. Nat. Neurosci. 18, 282–288.

Wu, X., and Foster, D.J. (2014). Hippocampal Replay Captures the Unique Topological Structure of a Novel Environment. J. Neurosci. 34, 6459–6469

Yang, K., and Dani, J.A. (2014). Dopamine D1 and D5 Receptors Modulate Spike Timing-Dependent Plasticity at Medial Perforant Path to Dentate Granule Cell Synapses. J. Neurosci. 34, 15888–15897.

Zhang, J.C., Lau, P.M., and Bi, G.Q. (2009). Gain in sensitivity and loss in temporal contrast of STDP by dopaminergic modulation at hippocampal synapses. Proc. Natl. Acad. Sci. 106, 13028–13033.

